# Isolated nuclei from frozen tissue are the superior source for single cell RNA-seq compared with whole cells

**DOI:** 10.1101/2023.02.19.529150

**Authors:** Andrew Jiang, Klaus Lehnert, Suzanne J. Reid, Renee R. Handley, Jessie C. Jacobsen, Syke R. Rudiger, Clive J. McLaughlan, Paul J. Verma, C. Simon Bawden, Russell G. Snell

## Abstract

The isolation of intact single cells from frozen tissue is a challenge due to the mechanical and physical stress inflicted upon the cell during the freeze-thaw process. Ruptured cells release ambient RNA into the cell suspension, which can become encapsulated into droplets during droplet based single cell RNA-seq library preparation methods. The presence of ambient RNA in droplets has been suggested to impact data quality, however there have been limited reports on single cell RNA-seq data from frozen tissue. Here, we compare the results of single cell RNA-seq derived from disaggregated cells from frozen brain tissue with single nuclei RNA-seq derived from purified nuclei of identical tissue using the 10X Genomics Chromium 3’gene expression assay. Our results indicated that presence of ambient RNA in the cell suspension resulted in single cell RNA-seq data with a 25-fold lower gene count, a 5-fold lower UMI count per cell and a 2-fold lower fraction of reads per cell compared with single nuclei RNA-seq data. Cell clustering with the single cell RNA-seq data was unable to resolve the heterogeneity of brain cell types. Our conclusion is that nuclei from frozen tissue are the superior substrate for single cell transcriptome analysis.

## Introduction

Obtaining fresh tissue is often not practical when conducting single cell transcriptome experiments. As a result, frozen tissue is typically the starting resource for single cell RNA-seq (scRNA-seq). Recovery of individual intact cells from frozen tissue is difficult due to the mechanical and physical stress inflicted upon the cell during the freeze-thaw process. The freeze-thaw process alone can cause cell membrane rupture and result in a release of ambient RNA into cell suspensions which can be partitioned into microfluidic droplets in the library making process. The co-encapsulation of ambient RNA with native cellular RNA can result in cross contamination of transcripts from multiple different cells and cell populations. This will have downstream consequences in clustering and can confound data analysis [1]. Therefore, scRNA-seq library preparation protocols recommend performing single nuclei RNA-seq (snRNA-seq) as an alternative to scRNA-seq for frozen samples [2, 3]. Nuclei are more resistant to damage incurred in the freeze-thaw process. Nevertheless, snRNA-seq presents other problems. First, lower amounts of mRNA are recovered from nuclei than whole cells and this results in RNA libraries with lower numbers of total genes per nuclei and unique molecular identifiers (UMIs) per nuclei which affects clustering [3]. Second, techniques that include quantification of cell surface markers (i.e., CITE-seq, 10X multiome scRNA-seq + ATAC-seq) cannot be performed on nuclei. Lastly, mitochondrial transcripts are completely absent in nuclei.

There have been limited reports of scRNA-seq experiments performed on frozen tissue and therefore our aim was to explore the performance of scRNA-seq on this tissue source. We compare the results of single cell RNA libraries prepared from post-mortem striatal (brain) tissue of four sheep against single nuclei RNA libraries prepared from identical tissue using the 10X Genomics Chromium 3’ gene expression assay.

## Methods

### Frozen brain tissue collection

The tissues used here were isolated from the striatum of four 5-year-old sheep. Tissue collection was performed as described in previous publications [4, 5] and performed in accordance with the SARDI/PIRSA Animal Ethics Committee (approvals no. 19/02 and 05/12). All experiments performed adhered to the recommendations in the ARRIVE guidelines. In brief, to obtain the sheep brains a lethal dose of pentobarbitone sodium solution (Lethabarb, 1 mL/2 kg body weight) was administrated intravenously, the brains were removed from the skulls, dissected into samples representing various brain regions and snap frozen initially on dry ice and then in liquid nitrogen. Animal brain weight, body weight and post-mortem delay is detailed in Table 1. Samples were wrapped in tinfoil and stored at −80°C until further use. Preparation of cells and nuclei for scRNA-seq and snRNA-seq respectively was performed on subsamples dissected from the same striatal tissue.

**Table 1.** Sheep cohort sampling details.

| Sheep ID | Sex | Age at death (years) | Brain weight (g) | Body weight (kg) | Post-mortem delay (min)<sup>1</sup> |
| --- | --- | --- | --- | --- | --- |
| 1 | F | 5 | 132.46 | 86.6 | 41 |
| 2 | M | 5 | 123.67 | 74.2 | 50 |
| 3 | F | 5 | 128.19 | 75.6 | 45 |
| 4 | M | 5 | 129.24 | 91.4 | 62 |

<sup>1</sup>Post-mortem delay described as the time from death until striatal tissue was dissected and frozen.

### Preparation of cells for single cell RNA-seq

Dissociation of neural tissue for scRNA-seq was performed using a modified protocol from 10X Genomics (10X Demonstrated Protocol, CG00055). In brief, 2 mL of 2 mg/mL Papain (Sigma) was added to 30 mg of tissue and incubated at 37 °C for 20 minutes. The papain solution was then removed without disrupting the tissue, leaving enough solution to cover the tissue. The tissue was dissociated using a Pasteur pipette with 2 mL of Neurobasal media (ThermoFisher Scientific), leaving cell debris to settle on the bottom. The supernatant containing the cells were transferred to a new 15 mL centrifuge tube and centrifuged at 200 xg for 2 minutes. The supernatant was removed, and the cell pellet resuspended in 1 mL Neurobasal media. 5 *μ*L of resuspended cells was stained with 5 *μ*L trypan blue to assess the cellular membrane intactness and number of cells was assessed using the Countess II FL Automated Cell Counter (ThermoFisher Scientific). The cell suspension was diluted by an appropriate volume of Neurobasal media to a final titer of 1000 cells/*μ*L. Cell suspensions from the tissues isolated from animals 1+2 and animals 2+3 were pooled resulting in a total of two cell suspensions.

### Preparation of nuclei for single nuclei RNA-seq

Nuclei were extracted from frozen ovine brain tissue utilising an adapted protocol from Krishnaswami et al., 2016 [2]. Briefly, approximately 50-100 mg of brain tissue was transferred to a dounce homogenizer containing 1mL homogenization buffer, HB (250mM sucrose, 25mM KCl, 5mM MgCl2, 10mM Tris buffer pH 8.0, 1μM DTT, 1X protease inhibitor, 0.4 U/*μ*L RNaseIn (ThermoFisher Scientific), 0.2 U/*μ*L Superasein (ThermoFisher Scientific), 0.1 % Triton X-100). Tissue was homogenised using 5 strokes with the loose dounce pestle A, followed by 10-15 strokes of tight dounce pestle B. The homogenate was filtered through a 40 *μ*m strainer into 5 mL Eppendorf tubes and centrifuged at 1000 xg (4 °C) for 8 minutes. The supernatant was removed and the pellet was resuspended in 250 *μ*L of HB. A 50% - 29% iodixanol gradient (OptiPrep™ Density Gradient Medium, Sigma) was prepared to allow removal of the myelin layer. 250 *μ*L of 50 % iodixanol was added to the nuclei-HB mixture and slowly layered on top of 500 *μ*L of 29 % iodixanol in a new Eppendorf tube. The resultant gradient was centrifuged at 13,000 g (4 °C) for 40 minutes. The supernatant and myelin was removed and the purified nuclei pellet was resuspended in 1 mL PBS, 1 % BSA, 0.2 U/*μ*l RNAse inhibitor. 5 *μ*L of resuspended nuclei was stained with 5 *μ*L trypan blue and the quality and number of nuclei was assessed using the Countess II FL Automated Cell Counter (ThermoFisher Scientific). Nuclei suspensions from animals 1+2 and animals 3+4 were pooled resulting in a total of two nuclei suspensions.

### Library preparation, sequencing, and quality assessment

Single cell and single nuclei RNA libraries were prepared using the Chromium Next GEM Single Cell 3’ Reagent Kit v3.1 (10x Genomics) as per manufacturer’s instructions. Single cell RNA and single nuclei RNA libraries were prepared with a targeted cell or nuclei recovery of 8,000. Concentration and quality of the synthesised cDNA were assessed using the Bioanalyzer High-Sensitivity DNA Kit (Agilent, 5067-4627). cDNA was successfully amplified from the suspension containing 8,000 nuclei however cDNA prepared from the suspension containing 8,000 cells was undetectable and therefore library preparation was discontinued at this step. Due to an estimated 10 % of cells in the suspension containing an intact cellular membrane, a second pair of cell suspensions was prepared with a targeted recovery of 80,000 cells to ensure sufficient cDNA could be recovered. cDNA was successfully amplified from these cell suspensions and thus adapter ligation was continued. Sequencing ready libraries were assessed using the Bioanalyzer and sequenced on HiSeq Xten platform to generate on average 417,947,486 (range: 377,815,731 – 527,195,602) 150 bp paired end reads. Alignment of reads was performed using the CellRanger v6.0.1 pipeline with STAR v 2.7.2a to the sheep Oar_rambouillet_v1.0 reference genome and annotation (Ensembl release 107). Library quality was assessed using CellRanger’s web summary report and the ICARUS single cell RNA-seq analysis web server [6]. ICARUS was used to plot unique gene counts per barcode and UMIs mapped per barcode for quality analysis.

### Dimensionality reduction, clustering and cell type annotation

Gene count matrices were normalised and scaled by a factor 10,000. Dimensionality reduction with principal component analysis was performed using a set of 2,000 top variable genes (Seurat::FindVariableFeatures) [7]. Clustering was performed with 15 principal components, k-nearest neighbour value of 20 and the Louvain clustering algorithm. Cell type annotation was performed by comparison of cluster marker genes against known striatal cell type markers identified in published scRNA-seq datasets [8–10].

## Results

Single cell and single nuclei RNA libraries were prepared using isolated cells or nuclei from frozen sheep striatal tissue. cDNA recovered from the two nuclei suspensions containing 8,000 nuclei showed a cDNA length distribution with maxima at approximately 1200 bp (Figure 2A). The shape of the pherogram for nuclei samples were consistent with 10X Genomics in-house testing for brain samples [11]. The initial attempt to construct a single cell RNA library targeted 8,000 cells. Microscopic examination of intact cellular membranes as screened by trypan blue viability exclusion assay showed 80% and 90% ruptured cells in the two cell suspensions. This fraction of ruptured/unruptured cells is unsurprising given the difficult nature of isolating intact cells from frozen tissue. The freeze-thaw and extraction process induces mechanical and physical stress to the cellular membranes leading to widespread cell lysis and release of mRNA [12, 13]. The cell pools containing both unruptured and ruptured cells were carried forward to cDNA preparation, but no cDNA was detectable in the bioanalyzer pherograms (Figure 1A); therefore library preparation from these samples was discontinued at this stage. A second pair of cell suspensions was prepared with a targeted recovery of 80,000 cells to ensure sufficient cDNA for the library making process. The percentage of ruptured cells in the second pair of suspensions was estimated at approximately 85% and 93% respectively. cDNA was detected in the Bioanalyzer pherograms (Figure 1A). Although cDNA yield was within the desired range, higher amounts of cDNA in the 350-600 bp range was observed that differed from the size distribution of cDNA from nuclei preparations. Following SPRIselect size selection, sequencing-ready libraries for both nuclei and cell samples had typical cDNA length distribution at 300-1000bp (Figure 1B & 2B).

**Figure 1.**
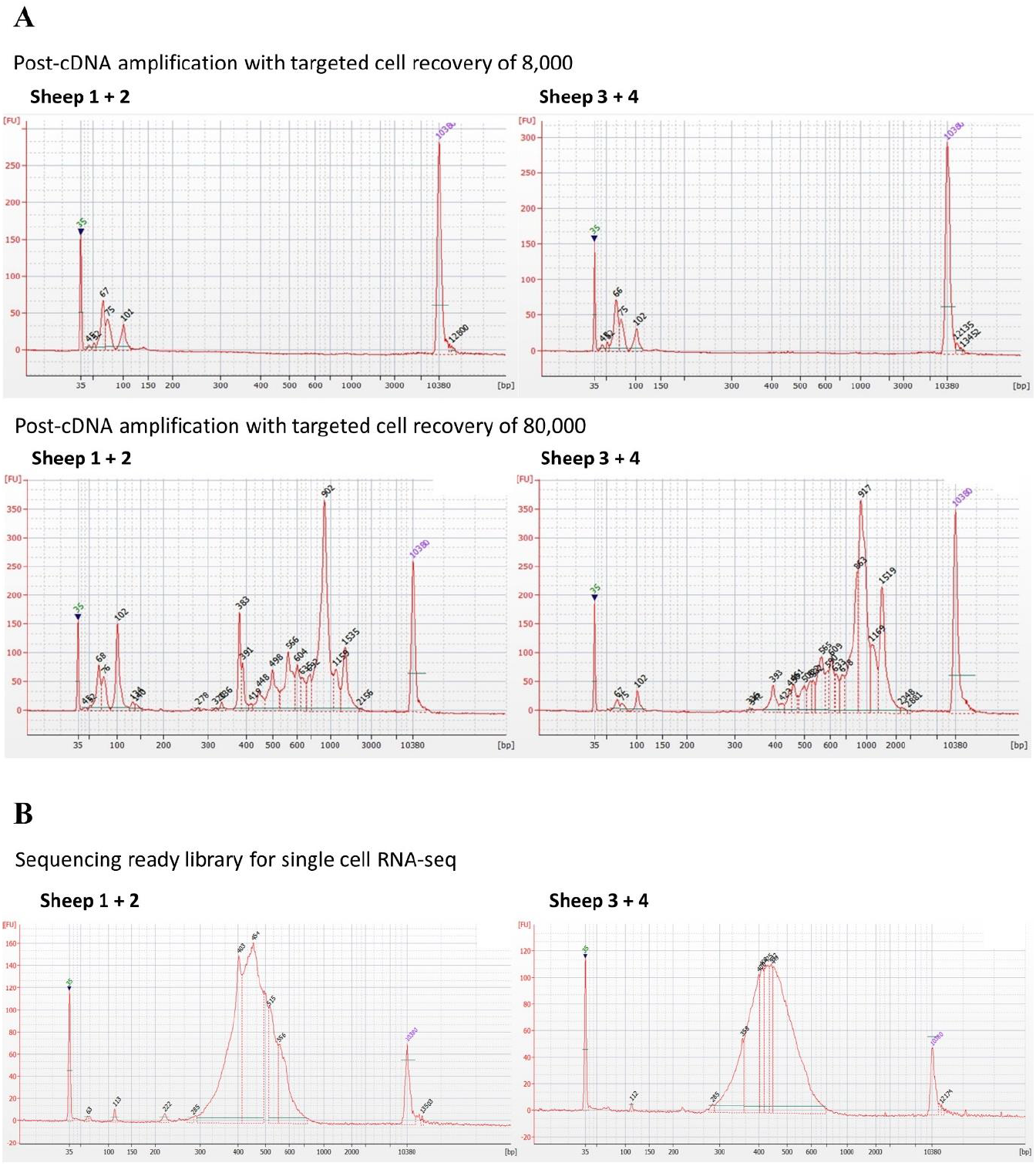
Bioanalyzer pherograms of single cell cDNA and sequencing ready libraries. (A) cDNA libraries with targeted cell recovery of 8,000 and 80,000 were prepared from frozen sheep striatal tissue using the 10X Chromium Next GEM Single Cell 3’ v3.1 Kit. No cDNA was detected for the cell suspensions containing 8,000 cells. Higher amounts of cDNA in the 350-600 bp range (indications of ambient RNA) were detected for the cell suspensions containing 80,000 cells. (B) Sequencing ready libraries exhibited a typical cDNA length distribution at 300-1000bp due to SPRIselect size selection.

**Figure 2.**
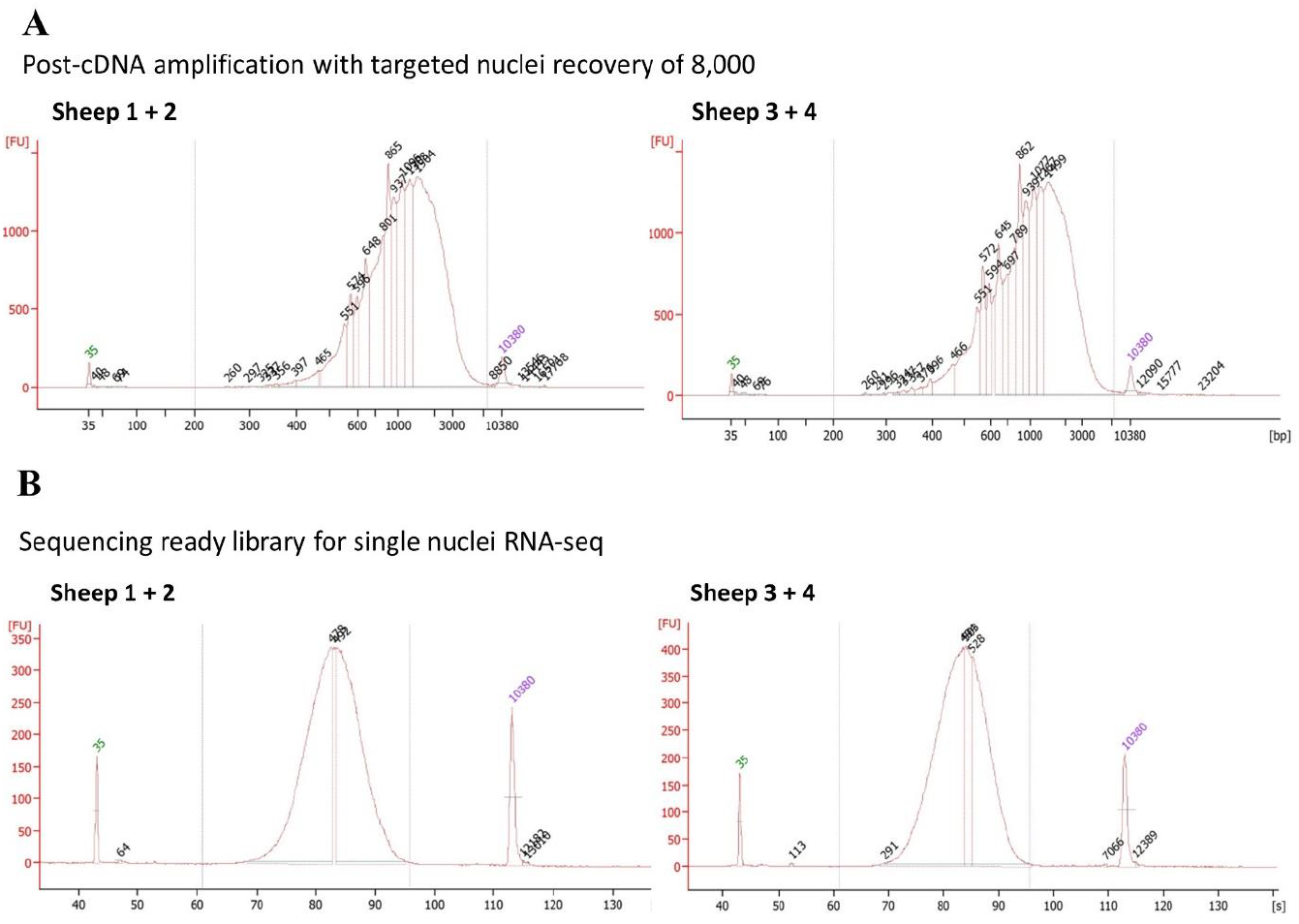
Bioanalyzer pherograms of single nuclei cDNA and sequencing ready libraries. (A) cDNA libraries and (B) sequence libraries were prepared from frozen sheep striatal tissue using the 10x Chromium Next GEM Single Cell 3’ v3.1 Kit. The shape of the pherogram was consistent with 10X Genomics in-house testing for brain tissue [8]. (B) Sequencing ready libraries exhibited a typical cDNA length distribution at 300-1000 bp due to SPRIselect size selection.

Reads were aligned to the Ovis aries reference genome with the 10X Genomics CellRanger software. CellRanger summaries indicated excellent read quality with >90% of barcodes, reads and UMIs containing a Phred score of Q30 or above for single cell and single nuclei RNA libraries (Table 2). 89.5% and 93.1% of reads from the single cell RNA libraries, and 96% and 98.9% of reads from the single nuclei RNA libraries, were successfully mapped to the reference genome. A ranked plot of every detected 10X barcode in decreasing order of number of UMIs associated with that barcode (barcode rank plot) is used to estimate the number of cells/nuclei in each library. The number of cells/nuclei is estimated by observing a distinctive steep dropoff at two inflection points, known as the cliff and knee (blue and red arrows respectively, Figure 5) that indicates a clear separation between barcodes associated with high UMI count (cell/nuclei associated barcodes) and barcodes associated with low/absent UMI counts (empty droplets). A characteristic cliff and knee were observed in single nuclei RNA libraries, whereas the steep cliff was not detectable in the data from the single cell RNA libraries. This has resulted in a substantially different number of estimated cells/nuclei between single cell and single nuclei RNA libraries. For the single nuclei RNA libraries, 8,815 and 8,723 barcodes were recovered. In contrast, for single cell RNA libraries, 2,162 barcodes were detected in one library and 32,500 barcodes for the other library. The fraction of reads in barcodes metric is the percentage of reads that are associated with a valid barcode and are mapped to the transcriptome. This fraction was low in single cell RNA libraries (17% and 60.8%) compared to single nuclei RNA libraries (90.3% and 78.4%). The fraction of reads in barcodes is usually >70% for a typical single cell RNA and single nuclei RNA libraries [14]. The single cell RNA libraries also contained very low UMI per cell (median UMI counts 348 and 504), and consequently low numbers of genes detected per cell (median gene count 40 and 72) (Figure 3). Both metrics were well below single nuclei RNA libraries (median UMI counts 2,612 and 2,042; genes/nuclei 1,331 and 1,158). Median UMI counts per cell or nucleus usually range from 1,000 – 6,000, while the typical number of genes identified per cell exceeds 1,000 in most cell types and samples [15]. The percentage of reads mapped to the mitochondrial genome was much higher in single cell RNA libraries (43.8% and 42.1%) than in single nuclei RNA libraries (1.3% and 0.6%) (Figure 3). For single nuclei RNA libraries, unsupervised clustering produced distinct clusters that captures the expected heterogeneity of striatal cell types (Figure 4). A total of 13 cell types were identified including oligodendrocytes (expressing marker genes MOG and PLP1), medium spiny neurons (RGS9, PDE10A), oligodendrocyte precursor cells (PCDH15, PDGFRA), microglia (CSF1R, CX3CR1), astrocytes (AQP4, GFAP), neuroblasts (DCX) and interneurons (SST, NPY) (Supplementary 1). In contrast, for single cell RNA libraries, the same clustering procedure resulted in a single large cluster that failed to resolve the cell types in the striatum (Figure 4).

**Table 2.** Summary statistics for single cell RNA and single nuclei RNA libraries.

| Sheep IDs | Cells |  | Nuclei |  |
| --- | --- | --- | --- | --- |
|  | 1+2 | 3+4 | 1+2 | 3+4 |
| **Cells/nuclei targeted** | 80,000 | 80,000 | 8,000 | 8,000 |
| **Estimated Cells/Nuclei** | 2,162 | 32,500 | 8,815 | 8,723 |
| **% Sequencing Saturation<sup>1</sup>** | 92.4 | 75.5 | 47.0 | 57.0 |
| **Total Reads** | 384,394,271 | 382,384,343 | 377,815,731 | 527,195,602 |
| **Total Genes detected** | 9,405 | 15,622 | 21,520 | 21,569 |
| **Median UMI/Barcode** | 348 | 504 | 2,612 | 2,042 |
| **Mean Reads/Barcode** | 177,796 | 11,766 | 42,861 | 60,437 |
| **Median Genes/Barcode** | 40 | 72 | 1,331 | 1,158 |
| **% Fraction Reads in Barcode** | 17.0 | 60.8 | 90.3 | 78.4 |
| **% Reads Mapped to Genome** | 89.5 | 93.1 | 96.0 | 98.9 |
| **% Reads Mapped to Transcriptome** | 24.4 | 34.7 | 40.0 | 29.0 |
| **% Reads Mapped to Exonic Regions** | 24.8 | 35.1 | 6.5 | 8.1 |
| **% Reads Mapped to Intronic Regions** | 1.9 | 2.6 | 53.4 | 56.8 |
| **% Reads Mapped to Intergenic Regions** | 7.7 | 10.6 | 30.2 | 28.2 |
| **% Reads to the Mitochondrial Genome** | 43.8 | 42.1 | 1.3 | 0.6 |
| **% Reads Mapped Antisense to Gene** | 0.1 | 0.1 | 19.8 | 35.7 |
| **% Q30 Bases in Barcode** | 97.2 | 97.2 | 97.0 | 97.0 |
| **% Q30 Bases in RNA Read** | 90.6 | 90.5 | 91.3 | 91.1 |
| **% Q30 Bases in UMI** | 97.3 | 97.3 | 96.9 | 97.1 |

<sup>1</sup>Sequencing saturation defined as the fraction of reads originating from an already-observed UMI.

**Figure 3.**
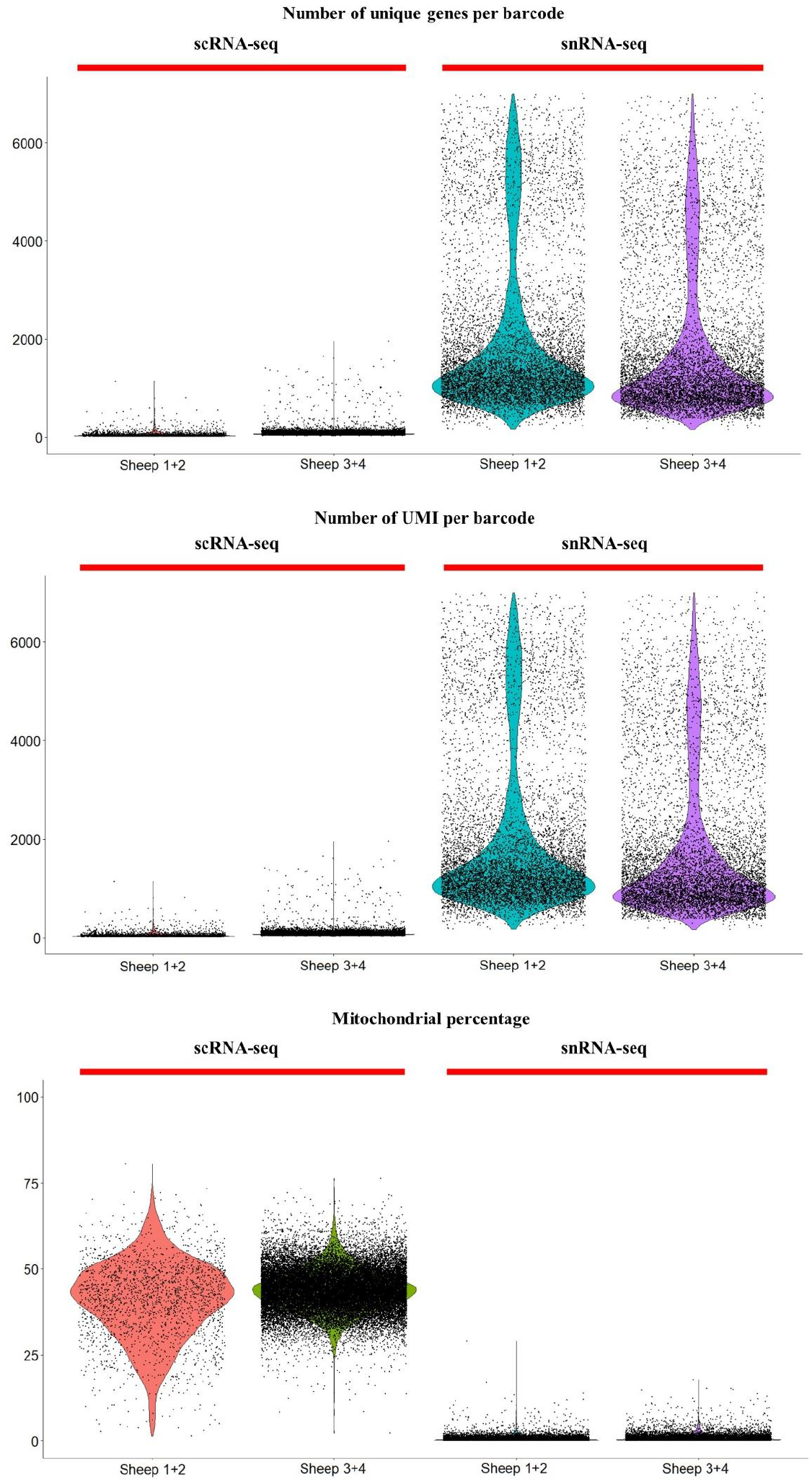
Quality control metrics for single cell RNA and single nuclei RNA libraries. Violin plots indicated single cell RNA libraries on average exhibited a 25-fold lower number of unique genes per barcode and a 5-fold lower UMI per barcode compared to single nuclei RNA libraries. The percentage of reads aligned to the mitochondrial genome (mitochondrial percentage) was also drastically higher in single cell RNA libraries.

**Figure 4.**
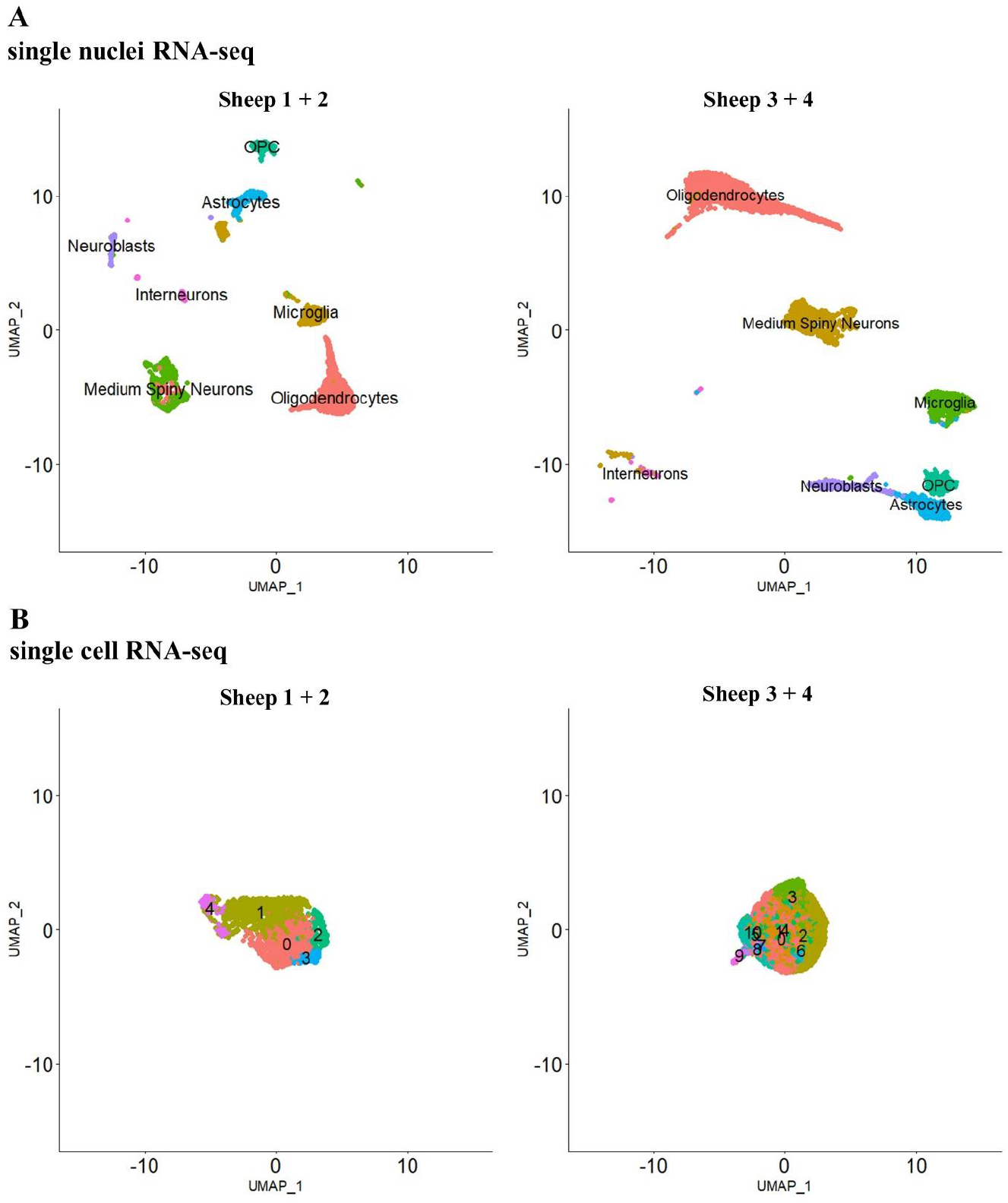
Clustering outcome for single cell RNA and single nuclei RNA libraries. (A) Single nuclei RNA-seq resulted in formation of distinct clusters representing the heterogeneity of gene expression for various cell types present within striatal brain tissue. (B) Single cell RNA-seq was unable to resolve the cellular heterogeneity of striatal cell types.

**Figure 5.**
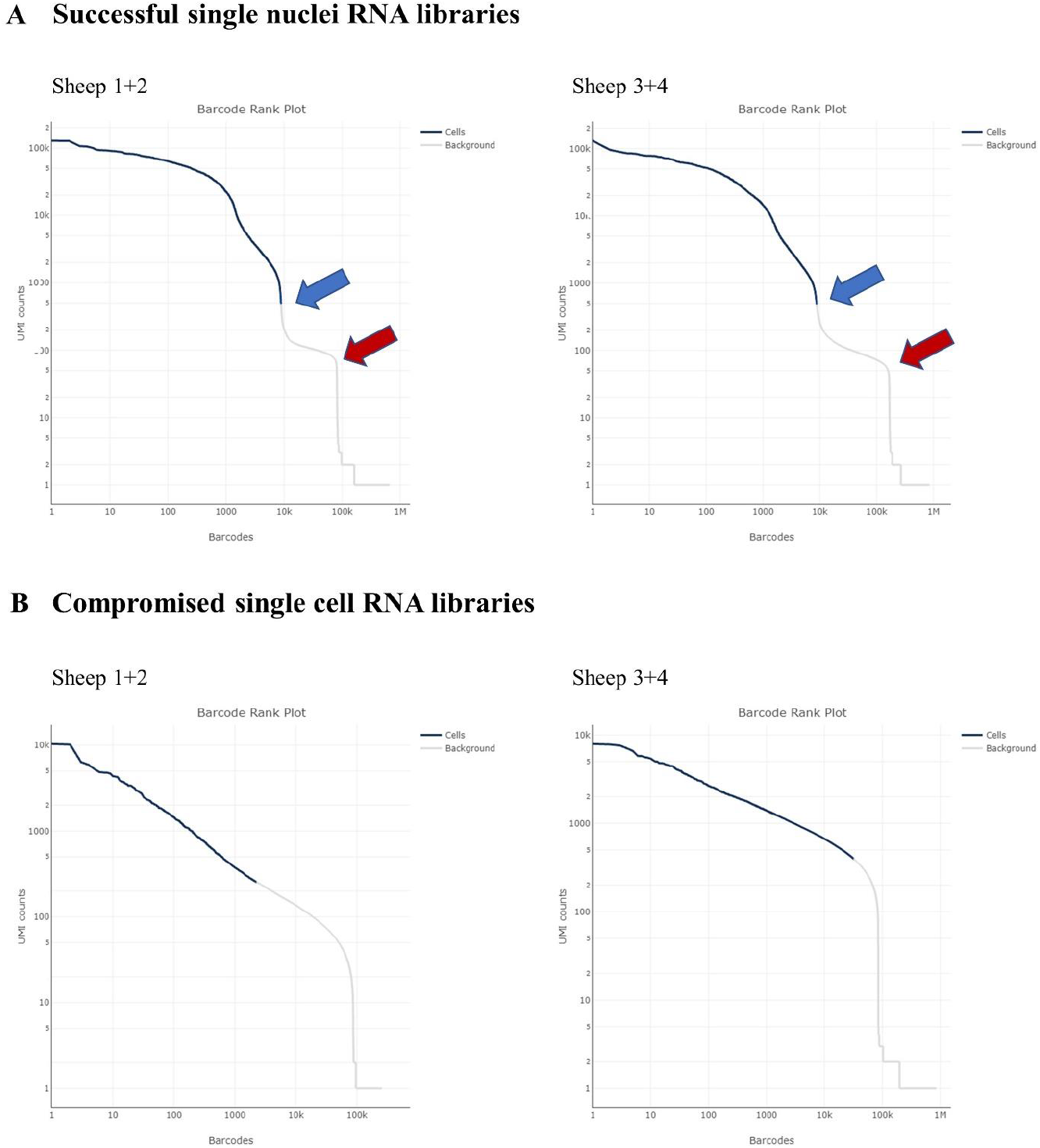
Comparison of barcode rank-plots from a successful single nuclei RNA-seq experiment versus compromised single cell RNA-seq experiments. (A) A barcode rank-plot from a successful experiment should exhibit a cliff (blue arrow) and knee (red arrow) shape indicating a clear separation between cell-associated barcodes and barcodes associated with low quality or empty GEMs. (B) A population of ambient RNA resulted in compromised barcode rank plots with a lack of steep cliff and no clear separation between cell-associated barcodes and empty GEMs.

## Discussion

This manuscript assessed the applicability of single cell RNA-seq for frozen brain tissue and compared the outcome of single cell RNA and single nuclei RNA libraries. We observed a very large proportion of cells to have ruptured membranes due to the freeze-thaw dissociation process (average ruptured cells estimated at 87%). These ruptured cells release RNA into the suspension (‘ambient RNA’) that can be partitioned with or without intact cells into a droplet gel bead (Gel Bead in Emulsion, GEMs). The ambient RNA is reverse transcribed to cDNA, which was detectable in bioanalyzer pherograms as increased amounts of short cDNA molecules in the 350-600 bp range. This was not evident in the nuclei derived samples. Further, libraries from single cell samples exhibited a compromised barcode rank-plot with a lack of steep cliff and no clear separation between cell-associated barcodes (high UMI counts) and empty GEMs. This lack of the anticipated graphical steep cliff is indicative of GEM encapsulation of cDNA from ambient RNA thereby diluting the number of empty GEMs. Consequently, a large disparity between the number of cell-associated barcodes detected in the single cell RNA libraries (2,162 and 32,500) was observed. The nuclei derived libraries in contrast revealed a consistent detection of nuclei associated barcodes (8,815 and 8,723) that conforms more closely to the initial targeted nuclei recovery of 8,000 per library. A higher proportion of reads that aligned to the mitochondrial genome was also observed in single cell RNA libraries. High mitochondrial content is often an indication of disrupted plasma membrane integrity. Mitochondrial transcripts are enclosed within a double membrane and are more resistant to membrane rupture. Hence, ruptured cells with intact mitochondrial DNA may be partitioned into GEMs, increasing the percentage of mitochondrial transcripts detected.

To address whether the presence of ambient RNA has affected data quality, we assessed common quality control metrics including the number of UMIs per barcode, number of unique genes per barcode and the fraction of reads in barcodes. Library quality of single cell RNA libraries was severely impacted shown by a 25-fold lower number of genes detected per barcode, a 5-fold lower number of UMIs attributed per barcode and a 2-fold lower fraction of reads attributed to a barcode compared to single nuclei RNA libraries. This has impacted cell clustering with clustering unable to resolve the cellular heterogeneity of striatal cell types. We conclude that maintaining plasma membrane integrity prior to library preparation is essential. Due to the difficulties in maintaining intact cells from frozen tissue, we recommend performing single nuclei RNA-seq as an alternative to scRNA-seq for frozen tissue. Nuclei are more resilient to the mechanical and physical stress induced by the freeze-thaw process and therefore allow the recovery of a high population of intact nuclei that represent the initial composition of the tissue. Our single nuclei RNA-seq datasets showcased a similar cell type composition and diversity to other single nuclei RNA-seq striatal datasets (Table 3) [10, 16]. Single nuclei RNA-seq has been widely applied to characterise transcriptomes from frozen brain tissue [17–24]. In summary, we have demonstrated successful application of single nuclei RNA-seq to frozen brain tissue that generates high quality transcriptomic data reflecting the transcriptome heterogeneity among cell types of the brain.

**Table 3.** Cell type composition of the striatum as determined by single nuclei RNA-seq datasets.

|  | Sheep 1+2 | Sheep 3+4 | Malaiya et al., 2021<br>[16] | Lee et al., 2020<br>[10] | Lee et al., 2020<br>[10] | Lee et al., 2020<br>[10] |
| --- | --- | --- | --- | --- | --- | --- |
| Species | Sheep | Sheep | Mouse | Mouse | Mouse | Human |
| <u>Cell types</u> |  |  |  |  |  |  |
| Medium Spiny neurons | 1,349 (15.3%) | 1,457 (16.7%) | 3,003 (66.4%) | 67,144 (61.9%) | 35,338 (59.3%) | 25,176 (20.1%) |
| Interneurons | 256 (2.9%) | 209 (2.4%) | 481 (10.6%) | 2,943 (2.7%) | 1,523 (2.5%) | 2,231 (1.8%) |
| Neuroblasts | 194 (2.2%) | 427 (4.9%) | NA<sup>1</sup> | 3,412 (3.1%) | 852 (1.4%) | NA<sup>1</sup> |
| Oligodendrocytes | 4,152 (47.1%) | 4,179 (47.9%) | 468 (10.3%) | 17,333 (16.0%) | 13,964 (22.9%) | 64,650 (51.6%) |
| OPC | 688 (7.8%) | 864 (9.9%) | 78 (1.7%) | 1,673 (1.5%) | 965 (1.6%) | 5,069 (4.0%) |
| Astrocytes | 811 (9.2%) | 654 (7.5%) | 300 (6.6%) | 11,696 (10.8%) | 5,185 (8.5%) | 14,819 (11.8%) |
| Microglia | 1,365 (15.5%) | 933 (10.7%) | 82 (1.8%) | 2,288 (2.1%) | 1,415 (2.3%) | 8,588 (6.9%) |
| Endothelial Cells | NA<sup>1</sup> | NA<sup>1</sup> | 112 (2.4%) | 343 (0.3%) | 186 (0.3%) | 902 (0.7%) |
| Mural Cells | NA<sup>1</sup> | NA<sup>1</sup> | NA<sup>1</sup> | 396 (0.4%) | 198 (0.3%) | 1,352 (1.0%) |
| Ependymal Cells | NA<sup>1</sup> | NA<sup>1</sup> | NA<sup>1</sup> | 1,239 (1.1%) | 470 (0.7%) | 2,384 (1.9%) |
| Total | 8,815 | 8,723 | 4,524 | 108,467 | 60,924 | 125,171 |

<sup>1</sup>Not described in dataset

## Acknowledgements

The authors wish to thank the incredible on-farm SARDI team for all animal management.

## Data Availability

The data have been deposited in NCBI’s Gene Expression Omnibus (GEO) [25] and are accessible through accession number GSE225915 (https://www.ncbi.nlm.nih.gov/geo/query/acc.cgi?acc=GSE225915).

## Author Contributions

A.J, K.L and R.G.S considered the experiment. S.J.R., R.R.H., S.R.R., C.J.M., collected the samples. A.J performed the single cell RNA-seq experiment and analysed the data. A.J., wrote the manuscript. K.L and R.G.S revised the manuscript. All authors reviewed and approved the final manuscript.

