## Supplementary material for "Isolated nuclei from frozen tissue are the superior source for single cell RNA-seq compared with whole cells": Supplmentary 1

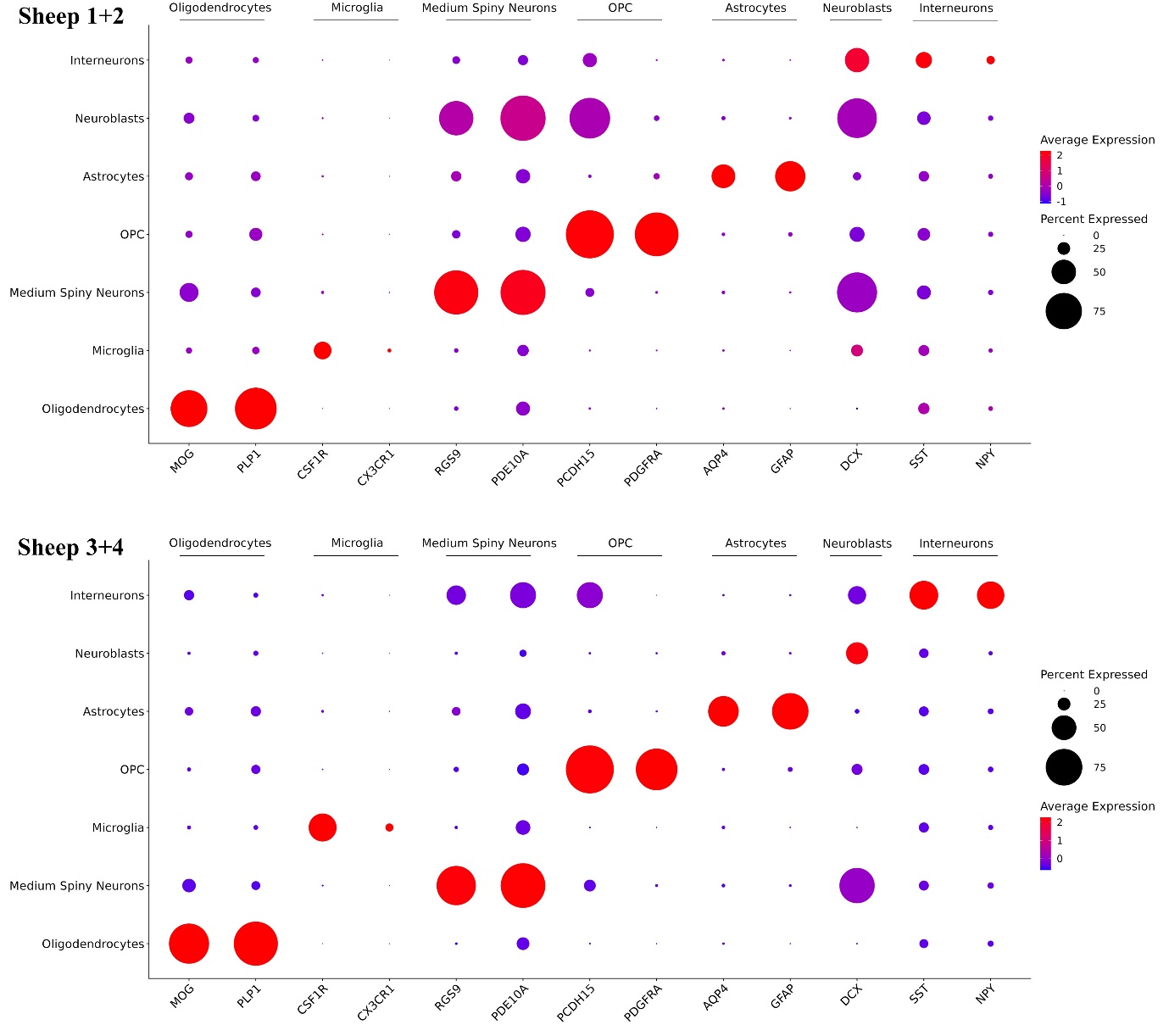
**Supplementary**

**Supplementary 1 Dot plot of enriched cell-type specific markers for annotated cell types in the single nuclei RNA-seq datasets.**
